# Generation of human induced trophoblast stem cells

**DOI:** 10.1101/2020.09.15.298257

**Authors:** Gaël Castel, Dimitri Meistermann, Betty Bretin, Julie Firmin, Justine Blin, Sophie Loubersac, Alexandre Bruneau, Simon Chevolleau, Stephanie Kilens, Caroline Chariau, Anne Gaignerie, Quentin Francheteau, Harunobu Kagawa, Eric Charpentier, Léa Flippe, Valentin Francois - - Campion, Sandra Haider, Bianca Dietrich, Martin Knöfler, Takahiro Arima, Jérémie Bourdon, Nicolas Rivron, Damien Masson, Thierry Fournier, Hiroaki Okae, Thomas Freour, Laurent David

## Abstract

Human trophoblast stem cells (hTSC) derived from blastocysts and first-trimester cytotrophoblasts offer an unprecedented opportunity to study the human placenta. However, access to human embryos and first trimester placentas is limited thus preventing the establishment of hTSC from a variety of genetic backgrounds associated with placental disorders. In the present study, we show that hTSC can be generated from numerous genetic backgrounds using post-natal cells *via* two alternative methods: (I) somatic cell reprogramming of adult fibroblasts using the Yamanaka factors, and (II) cell fate conversion of naive and extended pluripotent stem cells. The resulted induced and converted hTSC (hiTSC/hcTSC) recapitulated hallmarks of hTSC including long-term self-renewal, expression of specific transcription factors, transcriptome-side signature, and the potential to differentiate into syncytiotrophoblast and extravillous trophoblast cells. We also clarified the developmental stage of hTSC and show that these cells resemble post-implantation NR2F2+ cytotrophoblasts (day 8-10). Altogether, hTSC lines of diverse genetics origins open the possibility to model both placental development and diseases in a dish.

**Highlights:**

- Reprogramming of human somatic cells to induced hTSC with OSKM
- Conversion of naive and extended hPSC to hTSC
- Genetic diversity of hTSC lines
- Developmental matching of hTSC in the peri-implantation human embryo

## INTRODUCTION

During the first trimester of pregnancy, a subset of proliferative villous cytotrophoblasts (VCT) ensures the development and homeostasis of the placenta. These cells self-renew and differentiate into all cell types of the trophoblast lineage. Therefore, they are considered to be human trophoblast stem cells (hTSC).

Isolation of hTSC has been a major issue in the fields of developmental biology and stem cell research. Mouse TSC were derived in 1998, but hTSC were isolated only recently, in 2018, due to the difficulty to identify the compartment of these cells *in vivo* and the signaling pathways governing their self-renewal (Okae et al., 2018). Okae *et al* successfully derived hTSC from blastocysts and first trimester VCT. They designed a medium containing notably EGF, NODAL/TGFβ pathway inhibitors and a WNT pathway activator, which allowed prolonged culture of hTSC. In this manuscript, this medium is referred to as hTSC medium.

hTSC cultured *in vitro* represent a pristine model to investigate the development of the trophoblast lineage. These cells generate all differentiated trophoblast cell types, comprising the syncytiotrophoblast (ST) and extravillous trophoblasts (EVT) (Okae et al., 2018). The syncytiotrophoblast is the multinucleated outer layer of the trophoblast epithelium formed by cell-cell fusion of cytotrophoblasts. The unique structure of the ST facilitates diffusion of nutrients and gases between maternal blood and the fetus and protects the latter from pathogen entry. Extravillous trophoblasts (EVT) are migratory cells formed by epithelial-mesenchymal-like transition of cytotrophoblasts. EVT invade the decidual stroma, remodel the spiral arteries and participate to immune tolerance between the developing conceptus and the mother through a unique pattern of HLA expression, notably HLA-G (Knofler et al., 2019; Turco and Moffett, 2019).

hTSC at the origin of these processes play a central role in the formation of the maternal-fetal interface, and abnormal hTSC are likely to have dramatic consequences on placental development. This in turn can have postnatal outcomes and ultimately provoke chronic disease in the adulthood (Burton et al., 2016). However, we neither understand the nature and incidence of hTSC disorders, nor the connection of hTSC with placental diseases such as preeclampsia, fetal growth restriction, miscarriage or choriocarcinomas. For this purpose, researchers need to access hTSC of diverse genetic backgrounds associated with normal and pathological situations. So far, only few hTSC lines have been isolated from surplus embryos donated to research and aborted placentas, and we cannot access hTSC from individuals who were born after placental complications (Ezashi et al., 2019). To overcome these issues, we need alternative methods to generate hTSC from more accessible sources of cells.

Human induced Pluripotent stem cells (hiPSC), generated by somatic cell reprogramming, have the potential to differentiate into any cell type in the body and give access to patient specific cells (Kilens et al., 2018). Cell fate conversion is another method to generate cell types of interest, representing a faster approach which does not involve generation of iPSC lines. Chemical compounds can be sufficient to achieve cell fate conversion, which avoids transduction of exogenous factors (Kim et al., 2020). We hypothesized that these reprogramming strategies, largely applied to embryonic lineages, could also give access to extraembryonic cells, including those of the placenta. It has been reported that primed hPSC, corresponding to the post-implantation epiblast, acquire a trophoblast-like fate in response to BMP4, A83-01 (NODAL/TGFβ pathway inhibitor) and PD173074 (FGF pathway inhibitor), known as BAP treatment (Amita et al., 2013). However, these cells share features with differentiated trophoblasts and do not self-renew, which limits their use to model the human placenta. Also, the BAP model is debated, some claim that it produces mesoderm, others amnion-like cells (Bernardo et al., 2011; Guo et al., 2020).

In this study, we applied OSKM reprogramming of somatic cells and conversion of pluripotent stem cells to generate human induced and converted trophoblast stem cells (hiTSC and hcTSC), from patients with diverse genetic backgrounds. Comparison with isogenic hiPSC, placental cell types (VCT, VCT-ST, EVT), and previously-established hTSC lines confirmed that hi/cTSC share similar differentiation potential and molecular signature with embryo- and placenta-derived hTSC. This study paves the way to the production of patient specific hiTSC, with applications to obstetric medicine and the treatment of placental diseases.

## RESULTS

### Somatic cell reprogramming into hiTSC

We recently achieved reprogramming of somatic cells with OCT4, SOX2, KLF4, MYC (OSKM) into induced naive hPSC (hiNPSC), the counterpart of the preimplantation epiblast in human (Kilens et al., 2018). At low passage in t2iLGö medium (Takashima et al., 2014), we observed cobblestone-shaped morphology which was reminiscent of hTSC, in accordance with the expression of trophoblast associated transcription factor *GATA3* (**Figure S1A**). This suggested the occurrence of cells with dual potential to become either hTSC or hiNPSC, or a heterogeneous cell population. We thus investigated whether specific combinations of OSKM transgenes enabled to enrich the reprogrammed cells with hTSC. In the majority of cases, cells stopped proliferating at early passages. Only specific stoichiometries yielded naive hPSC but not hTSC lines (5:5:3 and 3:3:3 KOS, K, M multiplicity of infection). These results suggest that the stoichiometry of the OSKM reprogramming factors is not sufficient to reroute OSKM reprogramming towards trophoblast stem cells.

We thus hypothesized that the acquisition of the hTSC state might rather reside in environmental cues and repeated OSKM reprogramming in hTSC culture conditions. After 7 days, cells were transferred either in E7 medium that supports the early phase of reprogramming (Chan et al., 2009), or in hTSC medium. After 7 additional days, we observed the formation of epithelial colonies in both conditions, although these colonies were more abundant in E7. These results suggest that E7 not only supports the early phase of reprogramming but also promotes a mesenchymal-epithelial transition and the survival of reprogramming intermediates. Clearly, after culture in E7 (from day 7 to 21), cells robustly supported a transition into hTSC medium. Upon additional culture (2 passages), we observed the rapid formation of cobblestone-shaped colonies, morphologically reminiscent of hTSC (**Figure 1A**). These cell lines subsequently lost their transgenes (after 10 to 15 passages), and expressed the trophoblast associated genes *GATA2* and *GATA3*. In contrast, they did not express the pluripotency markers *NANOG* and *KLF17* (data not shown). These cells propagated unlimitedly, showing long-term self-renewal (> 70 passages). Based on their morphology, gene expression profile, and culture condition requirement, we referred to these reprogrammed cell lines as human induced trophoblast stem cells (hiTSC) (**Table S1A**).

**Figure 1.**
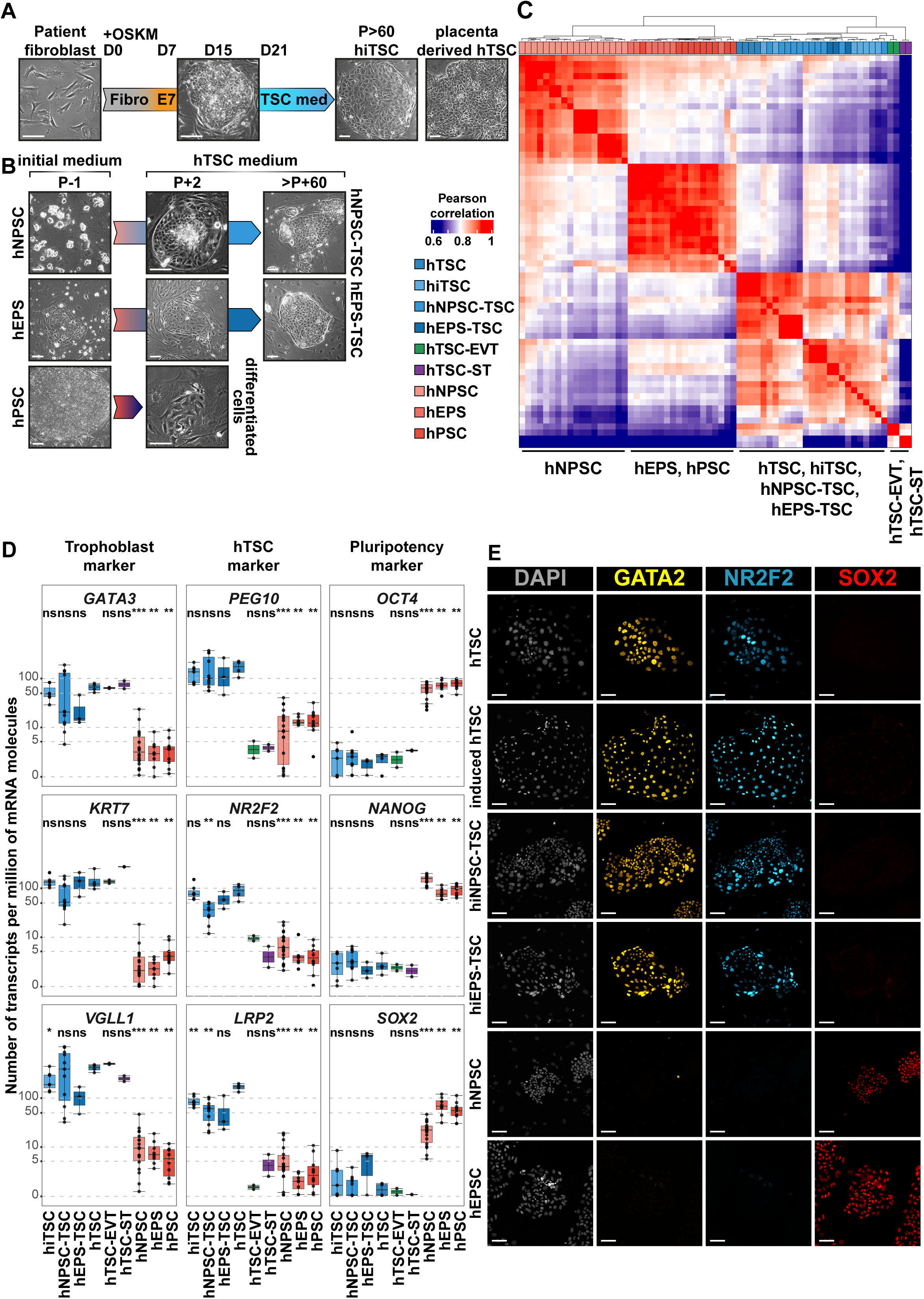
Generation of human induced trophoblast stem cells by reprogramming of somatic cells with OSKM and conversion of pluripotent stem cells. (A) Schematic representation of the reprogramming protocol. Phase contrast pictures show the changes in cell morphology. Placenta-derived hTSC are shown as controls. (B) Schematic representation of the conversion protocol. Phase contrast pictures show the changes in cell morphology. (C) Pearson correlation analysis of hPSC, hEPS, hNPSC, hiTSC, hcTSC and hTSC lines along with ST and EVT differentiated from hTSC. (D) Gene expression levels of indicated lineage markers are shown for hPSC, hEPS, hNPSC, hiTSC, hcTSC and previously-established embryo- and placenta-derived hTSC lines. The differentiated ST and EVT are included as controls. Expression levels are given as number of transcripts per million of mRNA molecules. A Wilcoxon-Mann-Whitney statistical test was performed for each type of hPSC and hTSC, with embryo- and placenta-derived hTSC taken as the reference group. Stars indicate statistical significance of the difference: *p-value < 0.05; **p-value < 0.01; ***p-value <0.001. (E) Immunofluorescence images of hTSC, hiTSC, hcTSC, hNPSC and hEPS stained for trophoblast-associated transcription factors GATA2 and NR2F2 and pluripotency associated transcription factor SOX2. Nuclei were stained with DAPI. Scale bars: 100 µm.

### Cell fate conversion of hNPSC and hEPS to hTSC

We tested the potential for primed hPSC that represent the post-implantation epiblast (Amit et al., 2000), extended hPSC (hEPS) that stabilize a high-potency state (Yang et al., 2017a), and naive hPSC (hNPSC) that reflect the pre-implantation epiblast (Kilens et al., 2018; Takashima et al., 2014) to respond to BAP treatment (Amita et al., 2013). Primed hPSC were initially cultured in KSR+FGF2 or iPS-BREW, hEPS in LCDM, and hNPSC in t2iLGöY medium. Consistent with previous reports, primed hPSC rapidly responded to BAP and transdifferentiated into large cell sheets morphologically reminiscent of trophoblasts. Interestingly, BAP culture could also induce similar morphological changes in hNPSC and hEPS, although these cell types are reflecting different states of pluripotency (**Figure S1B**). We confirmed the expression of trophoblast marker genes in all BAP treated hPSC, while we did not detect the expression of genes associated with other lineages, such as mesoderm or amnion (**Figure S1C**). However, these cells rapidly stopped proliferating and could not be maintained beyond day 16. We concluded that BAP medium efficiently promoted the conversion of hPSC to trophoblast-like cells independently from their initial state, but was not suitable for maintenance of self-renewing and expandable hTSC.

Then, we repeated the similar experiment in hTSC medium. Primed hPSC did not expand in the hTSC condition and we observed elevated cell death from 48h to 120h. Colony integrity faded after 1 or 2 passages and cells stopped proliferating thus failing to establish hTSC. Extended hPSC also experienced an elevated cell death from 48h to 120h, but few cobblestone-shaped colonies reminiscent of hTSC emerged after 2 passages (7-14 days) that could be expanded. In sharp contrast, naive hPSC sustained moderate cell death (48h-120h) and numerous colonies similar to hTSC rapidly emerged (7-14 days) and propagated unlimitedly (>50 passages, **Figure 1B**). These cells had lost expression of pluripotency markers *NANOG* and *KLF17* and gained expression of trophoblast associated genes *GATA2* and *GATA3* (data not shown). Hereafter, these cell lines are referred to as human converted trophoblast stem cells (hcTSC) (**Table S1A**).

### Molecular characterization of induced and converted hTSC

We conducted broad transcriptomic analyses to further characterize induced and converted hTSC in direct comparison with previously-established hTSC, and the differentiated ST and EVT (Okae et al., 2018).

Hierarchical clustering defined 3 groups of cells: (1) naive hPSC, (2) extended and primed hPSC, and (3) trophoblasts. Both induced and converted hTSC clustered together with previously established embryo-and placenta-derived hTSC to form the group of trophoblasts. This group subdivided between either hTSC comprising established, induced and converted hTSC ST and EVT. Pearson correlation analysis further confirmed the proximity of induced, converted, embryo- and placenta-derived hTSC. Surprisingly, extended and primed hPSC showed a high degree of transcriptional similarity, despite relative differences in their potential to form hTSC (**Figure 1C**). This can suggest that hEPS might contain rare subpopulations with higher potency comparable to hNPSC, or that the potential to form hTSC might rely on discrete cellular properties shared between hNPSC and hEPS.

Further analysis confirmed that induced and converted hTSC expressed key trophoblast lineage markers. Notably, the expression levels of *GATA3, KRT7* and *VGLL1* were similar to those found in previously established embryo- and placenta-derived hTSC (absolute gene expression ranging from 10 to 300 UPM). We also identified genes associated with stemness of hTSC, including *PEG10, NR2F2*, and *LRP2*. These were expressed at similar levels in hTSC, hiTSC and hcTSC, but not in the differentiated ST and EVT (absolute gene expression ranging from 20 to 200 UPM in hTSC, below 10 UPM in hTSC-ST and hTSC-EVT) (**Figure 1D**). In contrast, hiTSC and hcTSC did not express pluripotency associated markers such as *NANOG, SOX2* or *OCT4 (POU5F1)* (absolute gene expression below 10 UPM). We also confirmed that hi/cTSC did not express genes associated with other lineages (**Figure S1D**). Globally, gene expression profiles of hi/cTSC were comparable with those of embryo- and placenta-derived hTSC, but different from those of hPSC, which was confirmed by statistical analysis.

We finally analyzed hiTSC and hcTSC by immunofluorescence for the trophoblast markers NR2F2 and GATA2, that are expressed in the trophectoderm of human blastocysts (Meistermann et al., 2019a). NR2F2 and GATA2 were highly expressed and localized in nuclei of all cells. Conversely, SOX2 was highly expressed in hPSC, but absent in hTSC (**Figure 1E**). These expression patterns were comparable between hiTSC, hcTSC and placenta-derived hTSC. These results confirm that hi/cTSC share similar expression profiles with previously-established hTSC.

### Functional validation of hi/cTSC: differentiation into EVT and ST

hTSC are characterized by their ability to generate highly specialized trophoblast cell types that ensure the unique functions of the placenta. Notably, these cell lineages include ST and EVT. Therefore, we assessed the potential for induced and converted hTSC to differentiate into these cell types, using previously-established protocols based on NRG1 for EVT, and forskolin for ST differentiation (Okae et al., 2018). We directly compared this potential with those of embryo- and placenta-derived hTSC, VCT, ST and EVT cells isolated from human placentas.

Prior to differentiation assays, hTSC were cultured either in hypoxia or normoxia to promote EVT or ST formation, respectively (Chng et al., 2010; Wakeland et al., 2017). Initially, we cultured hTSC on MEF feeder layers that promoted undifferentiated proliferation as compared to other matrices. However, we observed that MEF impaired the formation of ST and EVT and therefore we screened for other matrices to adapt hTSC prior to differentiation assays. Fibronectin coatings were efficient at maintaining hTSC over time and laminin 521 could further enhance proliferation (**Figure S2A**). For an efficient differentiation, we adjusted the protocols of Okae *et al* as followed: EVT differentiation was enhanced by complementing the medium with IWR-1, which promoted the accumulation of EVT progenitors, as previously described ((Haider et al., 2018), pre-EVT medium). For ST and EVT differentiation, we adjusted cell density at seeding, from 0.8 to 6.0 × 10^4^ cells/cm^2^ and the timing of treatment initiation from 0h to 6h, depending on the lines (**Figure 2A-B**).

**Figure 2.**
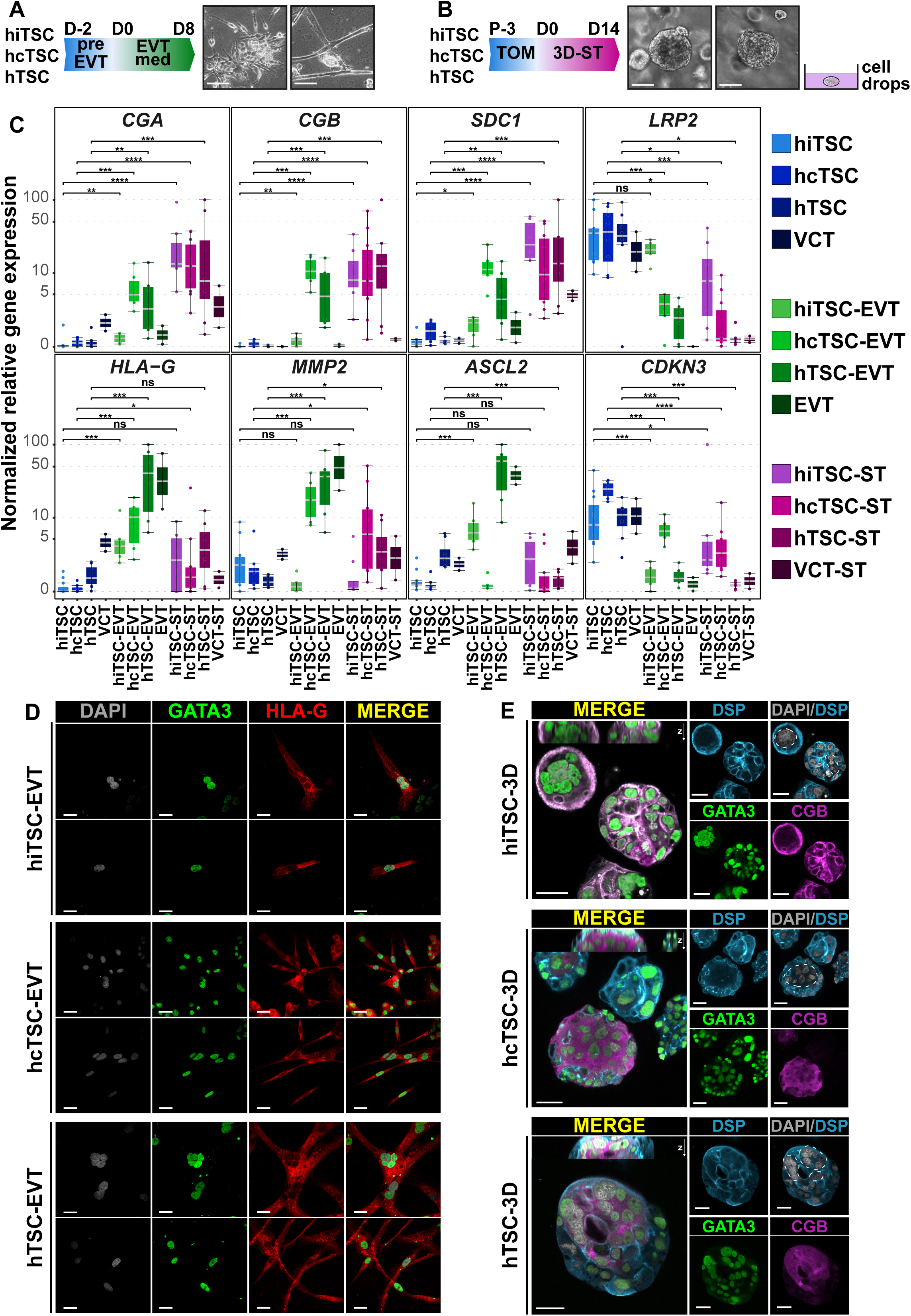
Induced and converted hTSC can differentiate into extravillous trophoblasts and the syncytiotrophoblast. (A) Schematic representation of the EVT differentiation protocol (left). Bright field pictures of the EVT progeny of h(i/c)TSC (right). (B) Schematic representation of the 3D-ST differentiation protocol (left). Bright field pictures of the 3D-ST structures derived from h(i/c)TSC (right). (C) RT-qPCR quantification of markers associated with ST (*CGA, CGB, SDC1*), EVT (*HLA-G, MMP2, ASCL2*) and hTSC (*LRP2, CDKN3*). A Wilcoxon-Mann-Whitney statistical test was performed for each type of hTSC and the differentiated cell progeny. Stars indicate statistical significance of the difference: *p-value < 0.05; **p-value < 0.01; ***p-value <0.001. (D) Immunofluorescence images of EVT differentiated from hiTSC, hcTSC and placenta-derived hTSC lines stained for the trophoblast-associated transcription factor GATA3 and the extravillous trophoblast-specific surface marker HLA-G. Nuclei were stained with DAPI. (E) Immunofluorescence images of 3D-ST structures derived from hiTSC, hcTSC and hTSC lines stained for GATA3 and the membrane-associated protein desmoplakin (DSP) highlighting syncytia, along with the syncytiotrophoblast-associated marker CGB. Nuclei were stained with DAPI. Scale bars (A-B): 100 µm; (D-E): 30 µm.

Upon optimized ST differentiation, cells upregulated the expression of *CGA, CGB* and *SDC1*, which are not expressed in hTSC (relative gene expression ranging from 10 to 1300-fold change). In contrast, *HLA-G, MMP2*, and *ASCL2* were globally increased in the EVT differentiation condition (relative gene expression ranging from 10 to 700-fold change). Finally, *LRP2* and *CDKN3* predominantly expressed in hTSC were downregulated upon differentiation (relative gene expression ranging from 2 to 70-fold change). Importantly, gene expression patterns were comparable with those of placental cells, which was confirmed by statistical analysis (**Figure 2C**).

Of note, some cell lines did not upregulate *MMP2* upon EVT differentiation, while others did not upregulate *ASCL2*. However, the clear expression of *HLA-G* unequivocally confirmed the EVT identity. This raises the possibility of distinct EVT populations, as previously described (Knöfler et al., 2019; Xiang et al., 2020) or might reflect intrinsic cellular properties of cell lines. In addition to specific gene expression, EVT can be reliably identified by dramatic morphological changes resulting in elongated shape (phase contrast images, **Figure 2A**). Immunostainings for GATA3 and HLA-G confirmed the identity of EVT differentiated from induced, converted and placenta-derived hTSC (**Figure 2D**).

In addition, ST can be identified by 2 important characteristics: the production of hCG and the fusion of cells that form multinucleated syncytia. After 6 days of forskolin treatment, β-hCG secretion was increased in h(i/c)TSC with a mean secretion superior to 3.9 × 10^3^ mIU/ml. In contrast, in those conditions, hPSC lines globally did not secrete β-hCG (mean expression inferior to 6.5 ×10^1^ mIU/ml). In line with transcriptomic analysis, both hPSC and hTSC lines secreted β-hCG when treated with BAP (**Figure S2D**).

Finally, it has been reported that the 3D culture of hTSC promotes the formation of ST (Haider et al., 2018; Okae et al., 2018). Based on trophoblast organoid formation protocols, we embedded hTSC as single cells in a semi-solid environment made of Matrigel, fibronectin and laminin 521. Cells were subsequently cultured in human trophoblast organoid medium (TOM) with small modifications (Turco et al., 2018a). Within a week, we observed the formation of 3D structures that grew to ∼200 µm in diameter after two weeks (**Figure 2B**). These structures contained multinucleated syncytia expressing DESMOPLAKIN (DSP) and CGB, and typically containing 6 to 10 nuclei (**Figure 2E**). In the majority of cases, fusion of cells occurred in the center of the 3D structures, as previously observed with placenta-derived trophoblast organoids (Haider et al., 2018; Turco et al., 2018a). Overtime, these placental organoids further grew until they reached confluency within the drops of matrigel. These larger structures also contained multinucleated syncytiotrophoblast cells (**Figure S2C**). Further optimization of culture conditions is needed to determine whether this system will allow expandable culture of hTSC-derived trophoblast organoids. We concluded that optimized differentiation protocols facilitate the formation of functional ST- and EVT-like cells and 3D self-organization from h(i/c)TSC lines. This confirmed the potential for induced and converted hTSC to form complex placental-like tissues including multiple differentiated and functional cell types.

### Dynamics of cell fate conversion from hPSC into hTSC

To further understand the conversion process of hPSC into hTSC, we projected samples on a PCA, which shows that samples cluster according to their fate. PC1 and PC2 accounted for 18% and 11% of variance, respectively. We identified 5 main clusters: (1) hNPSC, (2) hEPS/hPSC, (3) hTSC/hiTSC/hcTSC, (4) EVT, (5) ST (**Figure 3A**). PC3 accounted for 9% variance and segregated hNPSCs from other cells, thus confirming the particularity of naïve pluripotent stem cells, reflecting the early epiblast, and capable of efficiently generating hTSC (**Figure S3A-B**).

**Figure 3.**
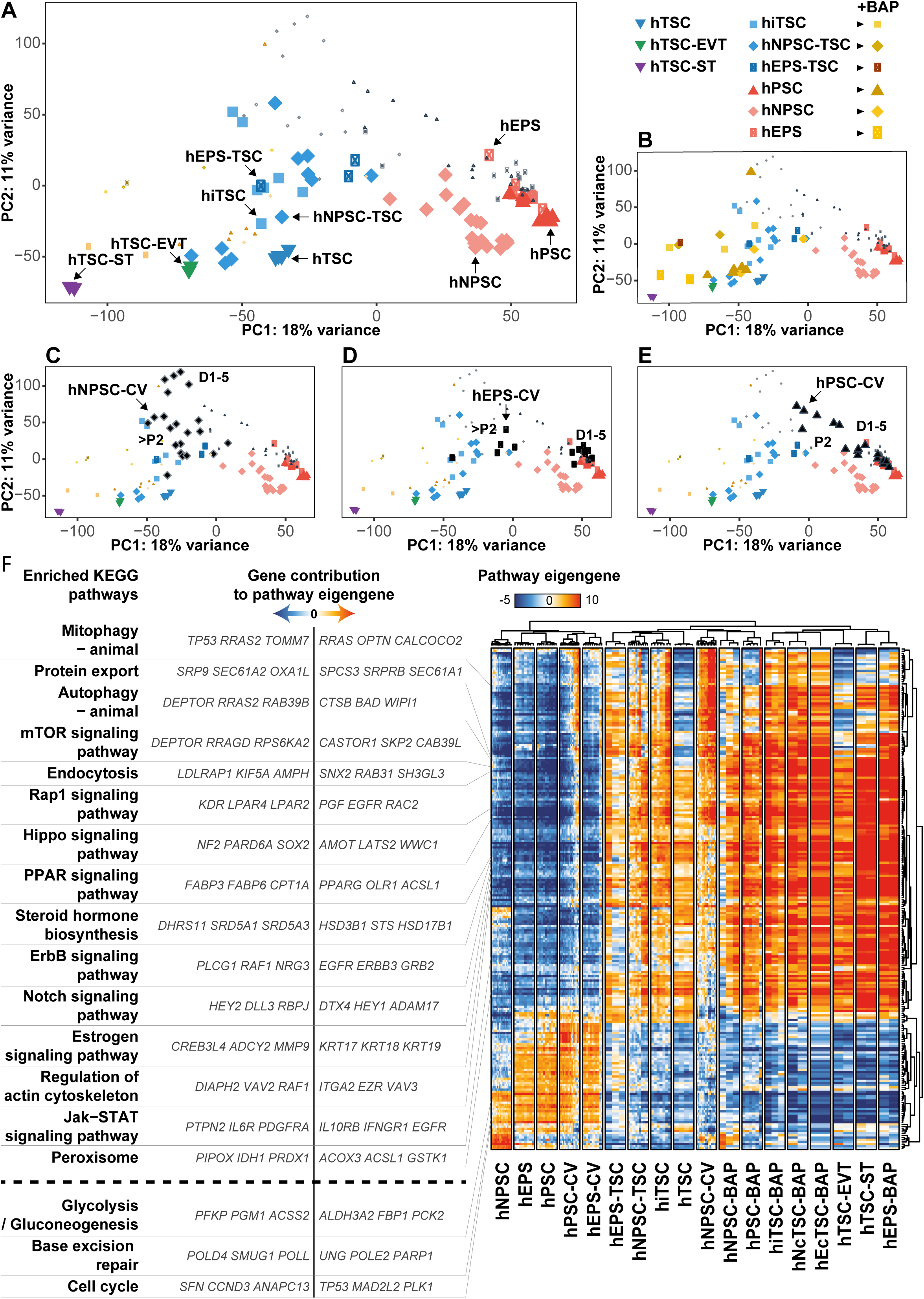
Transcriptional specificities of h(i/c)TSC and intermediate steps of cell fate conversion. (A-E) PCA analysis of h(i/c)TSC and hPSC in maintenance, differentiation or conversion media. BAP treated cells (B), naive hPSC converted into hTSC (C), extended hPSC converted into hTSC (D) or primed treated with hTSC media (E) are highlighted in specific panels. (F) Pathway enrichment analysis of h(i/c)TSC and hPSC in maintenance, differentiation or conversion media.

We further analyzed the progression of cells during cell fate conversion. All types of hPSC treated with BAP showed rapid and dramatic transcriptomic changes, and became close to differentiated trophoblasts by day 6. Despite some partial overlap with hTSC, they formed a distinct group, closer to the differentiated EVT and ST. Importantly, hTSC treated with BAP acquired a similar state, supporting that these transcriptional changes relate to the differentiation of the trophoblast lineage (**Figure 3B**).

Naive hPSC transferred to hTSC medium formed a separate cluster characterized by an intermediate transcriptome (P+2), followed by the acquisition of a hTSC molecular signature (∼P+3, **Figure 3C**). hEPS had delayed transcriptional variations and remained globally unchanged until day 5 before transiting through an intermediate transcriptional state (P+2), and ultimately acquiring a profile characteristic of hTSC (P+3, **Figure 3D**). We concluded that hNPSC and hEPS transited through an intermediate transcriptional state before ultimately achieving a cell fate conversion into hTSC. Whether this transcriptional progression reflects a developmental path remains to be investigated (Cinkornpumin et al., 2020). In contrast, primed hPSC initiated similar transcriptional changes but the cell fate conversion was not completed and cells did not acquire hTSC signature (P+2, **Figure 3E**).

Globally, treatments of hPSC with either BAP or hTSC medium converged to induce the acquisition of the trophoblast fate, but only hTSC culture condition supported the proliferation and self-renewal programs required to stabilize expandable stem cells. How external cues are integrated by cells to mediate these different outcomes will allow the identification of determinants of cell fate conversion, self-renewal, and proliferation of the native human placental progenitors.

### Modulations of signaling pathway signatures underly the conversion of hPSC to hTSC

We reasoned that signaling pathway variations upon conversion of hPSC to hTSC or differentiated trophoblasts might inform about the mechanisms of specification, self-renewal, and differentiation of human trophoblast progenitors. We thus performed a clustering analysis to identify co-regulated genes and a pathway enrichment analysis.

Clustering of samples on differentially regulated pathways globally confirmed the sample clustering previously performed on differentially expressed genes. This segregated 3 main groups: (1) hPSC, (2) hTSC, and (3) differentiated trophoblasts. In line with the PCA results, BAP-treated cells (day 6) were akin to ST and EVT and showed profiles of enriched pathways resembling those of differentiated trophoblasts. In contrast, hNPSC-TSC and hEPS-TSC shared similar pathway signatures with induced, embryo- and placenta-derived hTSC, while primed hPSC transited in hTSC medium globally failed to acquire hTSC pathway signatures (**Figure 3F**).

The clustering of pathways produced 2 main groups: (1) pathways associated with hPSC, and (2) pathways associated with trophoblasts. Pathways associated with hPSC included glycolysis, cell cycle and base excision repair, in line with previous studies on cell cycle regulation and metabolic state of hPSC (Kilens et al., 2018). In contrast, hTSC and differentiated trophoblasts globally shared common pathway signatures, clearly distinct from those of hPSC.

Differentially regulated pathways included HIPPO, NOTCH, and ERBB pathways, which are main drivers of the self-renewing state of mouse TSC (El-Hashash et al., 2010; Nishioka et al., 2009; Rayon et al., 2014; Rivron et al., 2018b), PPARG that is associated with mouse trophoblast proliferation and differentiation (Parast et al., 2009), and steroid hormone biosynthesis that is active in the human syncytiotrophoblast to produce placental hormones (Malassiné et al., 2003). Moreover, our analysis pointed to additional pathways including MTOR, Estrogen, RAP1 and JAK-STAT signaling pathways that seem to be active in h(i/c)TSC (**Figure 3F, S3C**).

Regarding the MTOR signaling pathway, genes with positive contribution to the eigengene were dominantly expressed in hTSC. They included *CASTOR1*, an inhibitor of mTORC1, and *PRR5*, a subunit of mTORC2. Also, trophoblast cells downregulated *RRAGD*, an activator of mTORC1. In contrast, hPSC expressed *MTOR, ULK1* and *SGK1*, which are effectors of mTORC1 signaling. These observations could indicate a switch from TORC1 in hPSC to TORC2 activity in hTSC during the cell fate conversion process. Interestingly, we also observed differences in MTOR pathway signatures between the distinct types of hPSC. Along with other genes, the expression of *DEPTOR*, a regulator of MTOR signaling, was high in hNPSC, moderate in hEPS and low in hPSC, suggesting a differential regulation of the pathway between these cells associated with different degrees of trophoblast potential (**Figure S3C**).

The estrogen signaling pathway seemed to be active in hTSC, which is reflecting the response to placental hormones that is observed *in vivo*. This was accompanied with the expression of *CREB3* and *FOS* that can mediate the transcription of estrogen responsive genes, and the upregulation of keratin genes, such as *KRT17, KRT18* and *KRT19*, in line with the morphological changes associated with the formation of epithelial hTSC (**Figure S3C**). Conversely to hTSC, hPSC expressed *FKBP5* and *HSP90AB1* which can form a complex that inhibits the translocation of the estrogen receptor to the nucleus (Baker et al., 2018).

hTSC also expressed *RRAS, VAV3* and *RAC2* which are effectors of the RAP1 signaling pathway, along with *EGFR* and *GNAI1*, which can signal to RAP1 through RTK and GPCR induced cascades, respectively. In contrast, hEPS and hPSC expressed *ID1* which is inhibited by RAP1, and globally did not express actors of the RAP1 signaling pathway, suggesting that it is not active in these cells. However, hNPSC expressed *RASGRP2* which specifically activates RAP1, along with *FGFR3* and *LPAR2* which encode for two receptors that can signal to RAP1 (**Figure S3C**). These gene expression profiles suggest that the RAP1 signaling pathway might be active in hTSC and hNPSC, but is mediated through different input signals, while it is not active in hEPS and hPSC.

We also found that hTSC expressed genes encoding receptors that can activate the JAK/STAT signaling pathway, including *IL10RB, CSF3R, CSF2RA* and *IFNGR1*. In contrast, hEPS and hPSC expressed *SOCS3*, a member of the suppressor of cytokine signaling protein family which inhibits JAK/STAT signaling, while hNPSC expressed *SOCS4* and *PTPN2*, which dephosphorylates JAK and STAT proteins thus inhibiting the signaling.

Interstingly, our analysis also pointed to gene expression profiles which suggested crosstalks between the identfied pathways. For example, *SFN* which is associated with cell cycle was specifically expressed in hTSC. When bound to KRT17, SFN regulates epithelial cell growth by stimulating the MTOR pathway (Kim et al., 2006). This suggests a potential cross-talk between the cell cycle, Estrogen and MTOR pathways in hTSC (**Figures 3F, S3C**).

Our analysis highlighted both conserved and divergent expression profiles of signaling pathways components underlying trophoblast specification and self-renewal. hTSC pathway signatures were globally milder than those of differentiated trophoblasts and the switch between the self-renewing state (hTSC) and the differentiated state (BAP treated cells) was mainly reflected by an accentuation of these same pathway signatures. This suggests that human trophoblast development is driven by a continuity rather than a sequential switch between different signaling activities, and that the strength of the signaling activity correlates with the progression from a self-renewal to a differentiation program. This dampened signaling activity observed in the self-renewing state might reflect the minimal requirement of unspecialized human trophoblast progenitors.

A comprehensive list of pathway components and their contribution to pathway eigengenes can be found in **Table S2G**. Further KO experiments are needed to determine how these pathways are modulated between hPSC, hTSC and trophoblast lineages, yet, this analysis gives a global picture of hTSC pathway signatures, and changes associated with cell fate conversion of hPSC.

### hTSC are akin to post-implantation day 8-10 cytotrophoblasts

An outstanding question regarding hTSC is to understand which developmental stage these cells are reflecting. To address this question, we compared the transcriptomes of hTSC with scRNAseq data of human peri-implantation embryos (146 embryos, 6838 cells), from day 3 to 14 (prolonged culture of human embryos for 9 days after blastocyst stage) (Petropoulos et al., 2016; Zhou et al., 2019). We projected all cells on a UMAP, and highlighted sample annotation (**Figure 4A**, *insert*). UMAP recapitulated developmental time and fate, showing the succession of eight cells, morula, early blastocyst, epiblast (EPI), primitive endoderm (PrE), trophectoderm (TE), and trophoblast (TB) cells from left to right. We noticed a cluster of trophoblast with enrichment of apoptosis-related genes, that we named “apoptotic trophoblast”, and the previously reported cluster of yolk-sac trophoblast (Zhou et al., 2019). We further clustered cells on the UMAP, which yielded 19 clusters of cells. In particular, this distinguished clusters associated with the development of the trophoblast lineage: early, medium and late pre-implantation TE, in line with our recent report (Meistermann et al., 2019b), 5 post-implantation trophoblast (TB #1 to TB #5), then pre-EVT, EVT, pre-ST and ST (**Figure 4A**).

**Figure 4.**
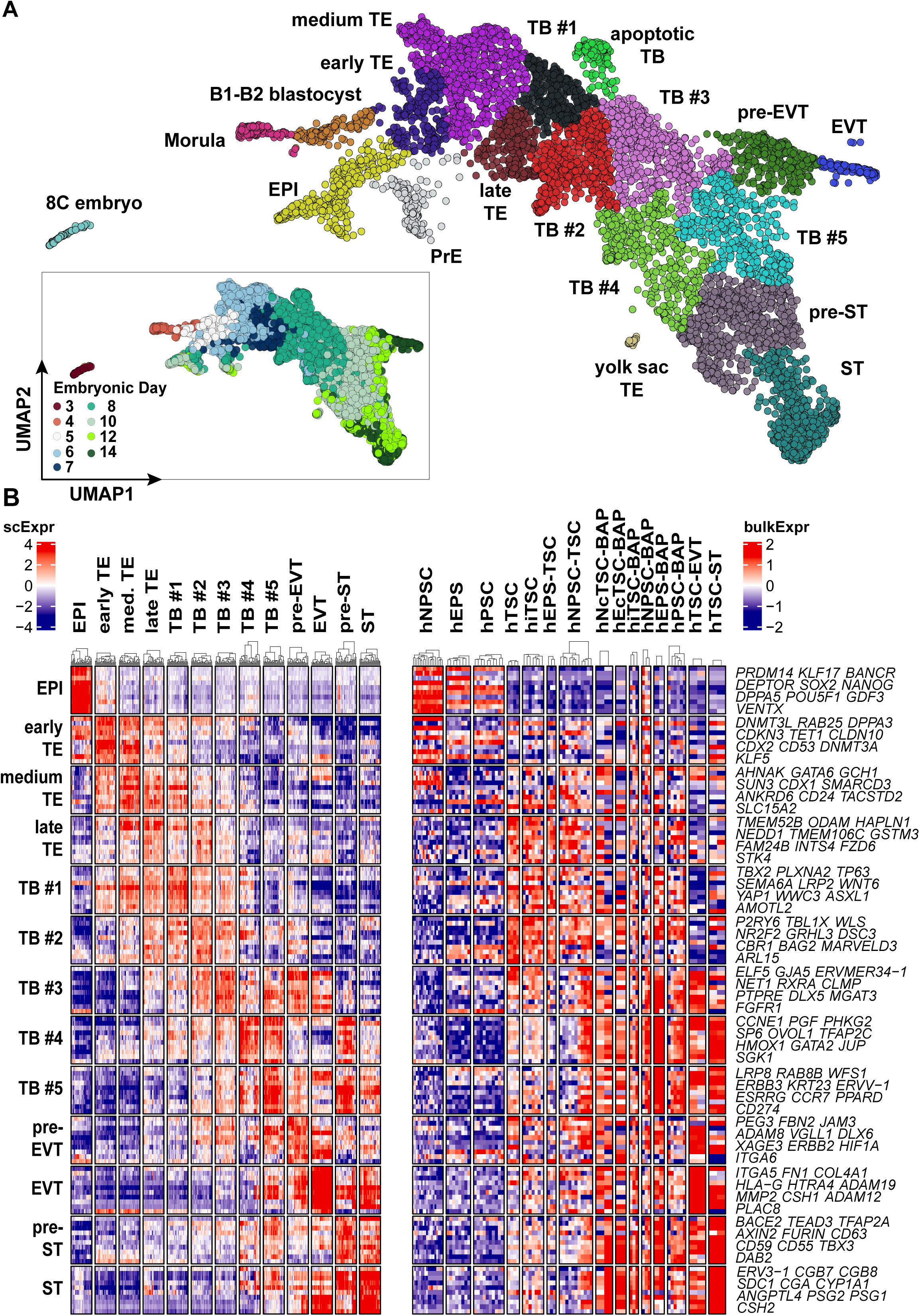
Developmental matching of hTSC with peri-implantation trophoblasts of the human embryo. (A) UMAP representation of scRNAseq datasets covering 8 cell stage to “day 14” of human development. Cluster analysis revealed 19 clusters, indicated on the UMAP. An insert shows the developmental day annotation on the UMAP. (B) Expression levels of selected genes characterizing clusters are shown for embryo (scRNAseq, left) and for h(i/c)TSC, hPSC and differentiated cells (DGEseq, right).

We used gene sets associated with each cluster, or gene sets recently associated with pre-implantation TE, post-implantation trophoblast, EVT and ST to benchmark the transcriptional signature of h(i/c)TSC (Xiang et al., 2020) (**Figure S4A-B**). Gene sets specific of each cluster highlighted the hierarchy of transcriptomic changes upon progression towards the trophoblast lineage (**Figure S4A**). Overall, comparison of transcriptional profiles of h(i/c)TSC with peri-implantation scRNAseq datasets pointed that h(i/c)TSC are related to the trophoblast lineage from day 5 to day 12 (**Figure S4A-B**). Curation of the list led us to propose markers of post-implantation trophoblast that matched hTSC lines, and distinguished them from ST, EVT and hPSC lines (**Figure 4B**). The markers that are better associated with hTSC are found in the gene sets specific of clusters TB #1, TB #2 and TB #3. Among those gene lists, we identified markers that have been assessed by immunofluorescence in human embryos: NR2F2, CDX2, GATA3, KRT7, CCR7 (Deglincerti et al., 2016; Meistermann et al., 2019b; Niakan and Eggan, 2013; Petropoulos et al., 2016). Projection of the expression of those markers on the UMAP led us to propose that hTSC expressed genes associated with clusters TB #1 and TB #2. Indeed, hTSC are expressing LRP2 and NR2F2, but not CDX2 (marker of medium TE) nor EVT or ST markers (**Figures 4B, S4C**). The clusters TB #1 and TB #2 are mainly composed of cells isolated from day 8 and day 10 embryos.

Altogether, our analysis points out markers that can distinguish developmental timing of TE and trophoblast cells, and associate hTSC with day 8-10 of development. Those markers notably include LRP2 and NR2F2.

## DISCUSSION

In the present study, we generated human induced trophoblast stem cells (hiTSC) from patients with diverse genetic backgrounds. We found that the OSKM system was not restricted to embryonic lineages, but was permissive to the trophoblast fate. Therefore, this system, largely accessible to researchers, is suitable for the parallel generation of isogenic hiTSC and hiPSC, which could greatly benefit to the study of placental diseases. Comparison with primary placental cells and hTSC lines confirmed that hi/cTSC were similar to embryo- and placenta-derived hTSC. They recapitulated transcriptome, protein markers and differentiation potential into EVT and ST.

In this study, we also revisited the relations between hPSC and the trophoblast lineage. We used two different systems to evaluate the potential of hPSC to generate hTSC: BAP treatment and transition into hTSC medium. We assessed a broad spectrum of hPSC, comprising: hNPSC, related to pre-implantation EPI, hEPS, showing contribution to extra-embryonic lineages in chimeras, and primed hPSC, related to post-implantation EPI. We found that all types of hPSC produced differentiated trophoblasts, but not hTSC, in response to BAP. In contrast, naive and extended hPSC, but not primed hPSC, converted to hTSC, following transition into hTSC medium.

During the preparation and revision of this manuscript, other groups have reported conflicting evidence that the potential to generate trophoblast is either higher in ground state or in expanded potential stem cells (Cinkornpumin et al., 2020; Dong et al., 2020; Gao et al., 2019). Gao et al claimed that expanded potential hPSC (hEPSC), but not naive hPSC (cultured in 5iLAF), can generate TSC. Dong et al, and Cinkornpumin et al reported conversion from 5iLAF naive cells. Here, we report successful conversion into hTSC from the two other main culture media used for naive hPSC (Takashima et al., 2014) and extended hPSC (Yang et al., 2017a). Our results indicate that the potential to engage the trophoblast lineage is common to all hPSC. However, in our model, only hNPSC and hEPS completed cell fate conversion to hTSC. We concluded that the potential to form hTSC correlates with the proposed developmental time equivalent of the initial culture, with naive hPSC being the most potent state to form hTSC. We do not exclude that primed hPSC could generate hTSC in another system, but this might rely on other pathways. Further investigations are needed to determine whether plasticity exists in the embryo, between the epiblast and the trophectoderm, and how it is regulated upon developmental progression.

Another key point of this study was to determine the developmental counterpart of hTSC in the embryo. To address this, we took advantage of single cell RNA-seq datasets of the human embryo during the peri-implantation development, from day 3 to 14. Our analysis revealed high complexity of the trophoblast lineage, divided in 9 clusters of cells, including TE, CTB, ST and EVT. We compared molecular signatures and found that hTSC resemble NR2F2+ day 8-10 CTB, which is clearly distinct from CDX2+ day 5-6 TE. Therefore, hTSC might be suitable to study early events of trophoblast lineage development, but maybe not pre-implantation TE. To address this, we need to optimize culture conditions to isolate and maintain human trophectoderm stem cells (hTESC). Recent studies suggest that this state exists in human, and CDX2+ hTSC have been obtained, but these cells have not been compared to the human TE yet (Knofler et al., 2019; Mischler et al., 2019).

Finally, generation of hiTSC by reprogramming provides a welcome alternative to the derivation of hTSC from embryos and placentas. This will enlarge the genetic repertoire of hTSC lines and give access to specific genetic backgrounds of interest. A next step is to generate hiTSC from patients affected by placental disorders. With this strategy, we can now consider studies to investigate the role of genetics in placental development and diseases such as preeclampsia, intrauterine growth restriction, miscarriage or choriocarcinomas. hiTSC could also serve for screening new formulations of human embryo culture media, with potential applications to IVF. New models of the embryo, such as blastoids, could benefit to this field of research (Rivron et al., 2018a). In this context, parallel derivation of isogenic hiTSC and hiPSC could provide a valuable source of cells. Overall, the assets of hiTSC are comparable to the advantages of hiPSC over hESC in the field of human development and disease modeling.

## ACKNOWLEDGMENTS

We thank the core facilities GenoBIRD, Micropicell, Cytocell, iPSCDTC. DM is funded by FINOX Forward Initiative. This work was supported by “Paris Scientifique region Pays de la Loire: HUMPLURI”.

## AUTHOR CONTRIBUTIONS

GC and BB performed experiments, with the help of other authors. JB performed biochemical dosages. DM, GC and SC performed bioinformatics analysis with the help of EC. GC and LD conceived the study and wrote the manuscript with the input of all authors.

## DECLARATION OF INTERESTS

DM is supported by FINOX forward grant initiative. Gaël CASTEL and Laurent DAVID have a provisional patent filled on the generation of human induced trophoblast stem cells.

## STAR METHODS

### EXPERIMENTAL MODEL AND SUBJECT DETAILS

#### Cell lines

For human somatic cell reprogramming into hiTSC and hiEPS, fibroblasts from healthy donors were used: BJ1, male fibroblasts are commercial BJ human neonatal fibroblasts extracted from normal human foreskin (Stemgent Cat# 08-0027); L71 from a 51-year-old healthy man; L80 from a 57-year-old healthy woman. Those fibroblasts were previously used to generate isogenic hPSCiPSC and hiNPSC (Kilens et al., 2018). For conversion experiments, we used hiNPSC and hiPSC lines from Kilens et al., and H9 hESC (WA09 Lot WB0090) obtained from the WiCell Research Institute. hESC were used under authorization RE17-007R from the French oversight committee, Agence de la Biomédecine. All cell lines used in this study are further described in Table S1A.

#### Human preimplantation embryos

Data analysis and transcriptomic modeling of human preimplantation development is detailed in Meistermann et al. (Meistermann et al., 2019a).

### METHOD DETAILS

#### Experimental design

Biological replicates are indicated in each figure. Randomization and blinding were performed for RT-qPCR but not for other experiments. No data or samples were excluded from any of the experiments.

#### Tissue culture

All cell lines were cultured at 37°C, under hypoxic (5% O_2_, 5% CO_2_) or normoxic conditions (20% O_2_, 5% CO_2_) as indicated. Culture medium was daily replaced. 10µM Y27632 (Axon Medchem™) was added to the culture medium upon cell seeding of human stem cells. PXX indicates passage number, and P+XX indicates that cells were converted for XX passages.

Human fibroblasts were cultured in fibroblast medium, composed of high glucose Dulbecco’s modified Eagle’s medium (DMEM) GlutaMAX-I (Gibco™) supplemented with 10% fetal bovine serum (FBS, Hyclone™), 1% sodium pyruvate (Gibco™) and 1% non-essential amino acids (Gibco™).

Mouse embryonic fibroblasts (MEFs) were prepared as previously described (Samavarchi-Tehrani et al., 2010) and cultured in fibroblast medium supplemented with 0.5% of penicillin–streptomycin (Life Technologies™). MEFs isolation was performed in compliance with the French law and under supervision of the UTE animal core facility, University of Nantes. MEFs were mitotically inactivated using mitomycin C to be used as feeder cells.

hiTSC were cultured on MEF feeder cells in hTSC medium [DMEM/F12 (Gibco™) supplemented with 0.1mM 2-mercaptoethanol (Gibco™), 0.2% FBS, 0.5% penicillin-streptomycin, 0.3% Bovine Serum Albumin (BSA, Sigma-Aldrich™), 1% Insulin-Transferrin-Selenium-Ethanolamine supplement (ITS-X, Gibco™), 1.5 mg/ml L-ascorbic acid (Sigma-Aldrich™), 50 ng/ml hEGF (Miltenyi Biotec™), 2 µM CHIR99021 (Axon Medchem™), 0.5 µM A83-01 (Tocris™), 1 µM SB431542 (Tocris™), 0.8 mM valproic acid (Sigma-Aldrich™) and 5 µM Y27632]. hiTSC could be passaged with TrypLE (15 min, 37°C, Life Technologies™) every 4 days at a 1:3 to 1:4 split ratio or every 7 days at a 1:40 to 1:60 split ratio. hiTSC were routinely cultured at 37°C in hypoxic conditions (5% O_2_, 5% C0_2_).

hNPSC were cultured on MEF feeder cells in t2iLGöY medium [DMEM/F12 supplemented with 1% N2 (Gibco™), 1% B27 (Gibco™), 1% non-essential amino acids, 1% GlutaMAX (Gibco™), 0.1 mM 2-mercaptoethanol, 50 µg/ml BSA, 0.5% penicillin–streptomycin, 1 µM CHIR99021, 1 µM PD0325901 (Axon Medchem™), 20 ng/ml mLIF (Miltenyi Biotec™), 5 µM Gö6983 (Axon Medchem™) and 10 µM Y27632] (Takashima et al., 2014). hNPSC were passaged every 4 days at a 1:3 split ratio using TrypLE (5 min, 37°C, Life Technologies™). hNPSC were routinely cultured at 37 °C in hypoxic conditions (5% O_2_, 5% CO_2_).

hEPS were cultured on MEF feeder cells in LCDM medium [48% DMEM/F12 and 48% Neurobasal (Gibco™) supplemented with 0.5% N2 supplement, 1% B27 supplement minus vitamin A (Gibco™), 1% non-essential amino acids, 0.1 mM 2-mercaptoethanol, 0.5% penicillin-streptomycin, 5% knockout serum replacement (KSR, Gibco™), 10 ng/ml human LIF (Miltenyi Biotec™), 1µM CHIR99021, 2 µM (S)-(+)-Dimethindene maleate (Tocris™) and 2 µM Minocycline hydrochloride (Tocris™), 1 μM IWR-endo-1 (Miltenyi Biotec™) and 2 µM Y-27632] (Yang et al., 2017b). hEPS were passaged every 4 days at a 1:8 split ratio using TrypLE (5 min, 37°C, Life Technologies™). hEPS were routinely cultured at 37°C in normoxic conditions (20% O_2_, 5% C0_2_).

Primed hPSC could be cultured on MEF feeder cells in KSR+FGF2 medium [DMEM/F12 supplemented with 20% KSR, 1% non-essential amino acids, 1% GlutaMAX, 50 µM 2-mercaptoethanol, 0.5% penicillin-streptomycin and 10 ng/ml human fibroblast growth factor 2 (FGF2, Peprotech™)]. 10 colonies were manually picked every 7 days for passage and seeded as small clumps (∼200 µm) in a new 35mm dish. Primed hPSC could also be cultured on MEF feeder cells in iPS-Brew medium (Miltenyi Biotec™) and these cells were passaged every 4 to 6 days at a 1:8 to 1:25 split ratio using TrypLE (5 min, 37°C, Life Technologies™). Primed hPSC were routinely cultured at 37°C in normoxic conditions (20% O_2_, 5% C0_2_).

10µM ROCK inhibitor (Y-27632) was systematically added to the culture media for 1 day after cell passaging with TrypLE. All cell lines were tested negative for mycoplasma using the MycoAlert kit (LONZA™, LT07-318).

#### Somatic cell reprogramming to hiTSC

Human adult fibroblasts were reprogrammed using the CytoTune-iPS 2.0 Sendai reprogramming kit (Life Technologies™). Two days before infection, 3.0 to 4.0 × 10^4^ fibroblasts were seeded per well on a 12-well plate, coated with Matrigel. At day 0, cells were infected with the three vectors: polycistronic Klf4-Oct4-Sox2, Myc and Klf4 at a 5:5:3 or 3:3:3 multiplicity of infection (MOI) respectively. At day 9 of infection, cells were dissociated with TrypLE (5 min, 37°C, Life Technologies™) and seeded in 35mm dishes, on MEFs. On the following day, cells were transited into E7 reprogramming medium (STEMCELL Technologies™). From day 21 onwards, cells were transited into hTSC medium. Induced hTSC lines (hiTSC) were routinely cultured at 37°C in hypoxic conditions (5% O_2_, 5%CO_2_).

Somatic cell reprogramming to hiNPSC, hiEPS and hiPSC was performed as described in previous reports (Kilens et al., 2018; Yang et al., 2017b).

#### Conversion of hNPSC and hEPS to hcTSC

hNPSC and hEPS were dissociated with TrypLE (5 min, 37°C, Life Technologies™) and seeded in 35mm dishes, on MEFs, at a density of 0.6 to 1.7 × 10^5^ cells per dish. Cells were maintained in their initial medium supplemented with 10 µM Y27632 for 1 day. From day 2 onwards, cells were transited into hTSC medium. Converted hTSC lines (hcTSC) were routinely cultured at 37°C in hypoxic conditions (5% O_2_, 5% CO_2_).

Primed hPSC included in conversion experiments were initially cultured in KSR+FGF2 or iPS-BREW. 10 colonies were picked (KSR+FGF2) or cells were passaged with TrypLE (iPS-BREW) and seeded at a density of 0.5 to 1.25 × 10^5^ cells per dish in 35mm dishes coated with MEFs for conversion assays.

#### Differentiation of hi/cTSC to EVT and ST

After at least 15 passages, cells were collected for differentiation assays. Prior to differentiation into ST and EVT, h(i/c)TSC (initially cultured on MEFs) were transited to fibronectin for at least 3 passages.

##### EVT differentiation

2-4 days before passage, h(i/c)TSC were transited into pre-EVT medium [DMEM/F12 supplemented with 0.1mM 2-mercaptoethanol, 0.5% penicillin-streptomycin, 0.3% BSA, 1% ITS-X supplement, 4% KSR, 7.5 µM A83-01 (Tocris™), 2.5 µM Y27632, 5 µM IWR-endo-1 (Miltenyi Biotec™)]. Then, cells were passaged with TrypLE to a density of 0.8 to 3.0 × 10^4^ cells/cm^2^. Before treatment was initiated, cells were placed in differentiation basal medium [DMEM/F12 containing 0.1 mM 2-mercaptoethanol, 0.5% Penicillin-Streptomycin, 0.3% BSA, 1% ITS-X], supplemented with 10 µM ROCK inhibitor (Y27632). Within 6 hours, timing depending on the lines, cells were transited into EVT medium (Okae et al., 2018) [Differentiation basal medium, supplemented with 100 ng/ml NRG1, 7.5 mM A83-01, 2.5 mM Y27632, 4% KnockOut Serum Replacement, 2% Matrigel]. At day 3, medium was replaced with the EVT medium containing 0.5% Matrigel, without NRG1. Typically, EVT formation was observed by day 4-5. At day 6, medium was replaced with the EVT medium containing 0.5% Matrigel, without NRG1 and KSR. Cells were collected on day 8 for subsequent analyses.

##### 2D-ST differentiation

h(i/c)TSC were passaged with TrypLE to a density of 2.0 to 6.0 × 10^4^ cells/cm^2^. Before treatment was initiated, cells were placed in differentiation basal medium [DMEM/F12 containing 0.1 mM 2-mercaptoethanol, 0.5% Penicillin-Streptomycin, 0.3% BSA, 1% ITS-X], supplemented with 10 µM ROCK inhibitor (Y27632). Within 3 hours, timing depending on the lines, cells were transited into ST medium (Okae et al., 2018) [Differentiation basal medium, supplemented with 2.5 mM Y27632, 2 mM forskolin, and 4% KSR]. Medium was replaced at day 3, and cells were analyzed at day 6.

##### 3D-ST differentiation

Prior to 3D differentiation assay, h(i/c)TSC were transited to trophoblast organoid medium (TOM) with small modifications (Turco et al., 2018b) [DMEM/F12, 1X N2 supplement, 1X B27 supplement minus vitamin A, 1.25 mM N-Acetyl-L-cysteine, 1% GlutaMAX (Gibco™), 0.5% Penicillin-Streptomycin (TOM basal medium), supplemented with 500 nM A83-01, 1.5 μM CHIR99021, 80 ng/ml human R-spondin1, 50 ng/ml hEGF, 100 ng/ml hFGF2, 50 ng/ml hHGF, 2 μM Y-27632]. 0.4 to 1.0 × 10^5^ cells passaged with TrypLE were embedded into 150µl drops comprising: 50µl Matrigel and 50µl PBS+/+ along with 960 ng fibronectin, 50 ng laminin521 and 50µl TOM basal medium. Drops were carefully deposited on sterile parafilm covered dishes and placed at 37°C for 20 minutes to solidify. Complete TOM supplemented with 10 µM ROCK inhibitor (Y27632) was further added to cover the drops. Medium was replaced every 3 days with TOM. The 3D structures emerged within a week and were collected on day14 for subsequent analyses.

#### MEF-BAP treatment

hPSC and hi/cTSC lines were dissociated with TrypLE and seeded on 35mm dishes coated with MEFs, at a density of 0.4 to 1.0 × 10^5^ cells per dish. For primed hPSC cultured in KSR +FGF2 medium, about 10 colonies were manually picked and seeded per dish. hPSC and hi/cTSC were maintained in their initial medium supplemented with 10 µM Y27632 for 1 day. The following day, initial medium was replaced with MEF-BAP [DMEM/F12 supplemented with 20% KSR, 1% non-essential amino acids, 1% GlutaMAX, 50µM 2-mercaptoethanol (MEF-CM, conditioned for 24 hours on a MEF monolayer and filtrated with a 0.22 µm pore size filtration unit), 2 ng/ml human BMP4 (Miltenyi Biotec™), 1µM A83-01 and 0.1 µM PD173074 (Axon Medchem™)]. MEFBAP medium was daily replaced. Supernatants and cells were collected at day 3 and 6 of differentiation for subsequent analyses.

#### Isolation of VCT, ST and EVT from human placentas

Isolation of placental cell types was conducted following previously published protocols (Handschuh et al., 2006). Briefly, chorionic villi were dissected from term placentas of healthy mothers. Mononucleated cytotrophoblasts (VCT) and extravillous trophoblasts (EVT) were isolated after trypsin-DNase digestion, sedimentation, filtration and discontinuous Percoll gradient fractionation. VCT were cultured for 2h in Dulbecco’s modified Eagle’s medium (DMEM) containing 10% decomplemented fetal calf serum (FCS), 2 mM glutamin, 100 IU/mL penicillin and 100 mg/mL streptomycin. EVT were cultured on Matrigel in the same culture conditions for 48h. ST were obtained by further differentiation of VCT for 72h in these conditions (spontaneous aggregation and fusion of VCT).

#### β-hCG dosage

Cell culture supernatants were collected at days 0, 3 and 6 of forskolin and MEF-BAP treatments. Amounts of secreted β-hCG were measured by electrochemiluminescence immunoassay “ECLIA” on Cobas® e601 immunoassay analyzer, at the Clinical Biochemistry Laboratory, CHU Nantes, France.

#### Immunostaining

For immunofluorescence (IF) analysis, cells were fixed at room temperature using 4% paraformaldehyde for 15 min. Samples were then permeabilized for 60 min at room temperature with IF buffer [phosphate-buffered saline (PBS), 0.2% Triton, 10% FBS], which also served as a blocking solution. Samples were incubated with primary antibodies overnight at 4 °C. The following antibodies were used: anti-GATA2 (1:50; Sigma® WH0002624M1), anti-GATA3 (1:100, R&D® AF2605), anti-NR2F2 (1:100, Abcam® ab211776), anti-CGB (1:100, Abcam® ab9582), anti-HLA-G (1:100, Abcam® 52455), anti-DSP (1:200, ref: Abcam® ab71690), anti-SOX2 (1:500, SCBT® sc-17320). Incubation with secondary antibodies was performed for 2 h at room temperature along with 4′,6-diamidino-2-phenylindole (DAPI) nuclei staining. Confocal immunofluorescence images were acquired with A1-SIM Nikon® confocal microscope. Optical sections of 0.5-1 µm-thick were collected. Images were processed using Volocity® visualization software.

#### RNA extraction and RT-qPCR

Total RNA was extracted using RNeasy® columns and DNAse-treated using RNase-free DNase (Qiagen). First-strand cDNAs were generated using 500ng of RNA, M-MLV reverse transcriptase (Life technologies), 25µg/ml polydT (Ambion) and 9.6µg/ml random primers (Roche). RT-qPCR were performed on a StepOne instrument (Applied Biosystems) using power SYBR green PCR master mix, for genes listed in the primers table (**Table S1B**). For each sample, the ratio of specific mRNA level relative to *GAPDH* levels was calculated. Experimental results are shown as relative gene expression.

#### Expression profiling by DGE-seq

For 3′ DGE, RNA-sequencing protocol was performed according to our implementation of M. Soumillon et al. protocol (Kilens et al., 2018; Magali Soumillon, 2014). Briefly, the libraries were prepared from 10 ng of total RNA. The mRNA poly(A) tails were tagged with universal adapters, well-specific barcodes and unique molecular identifiers (UMIs) during template-switching reverse transcription. Barcoded cDNAs from multiple samples were then pooled, amplified and tagmented using a transposon-fragmentation approach which enriches for 3′ends of cDNAs. A library of 350–800 bp was run on an Illumina HiSeq 2500 using a Hiseq Rapid SBS Kit v2-50 cycles and a Hiseq Rapid PE Cluster Kit v2. Read pairs used for analysis matched the following criteria: all 16 bases of the first read had quality scores of at least 10 and the first 6 bases correspond exactly to a designed well-specific barcode. The second reads were aligned to RefSeq human mRNA sequences (hg19) using bwa version 0.7.17. Reads mapping to several transcripts of different genes or containing more than 3 mismatches with the reference sequences were filtered out from the analysis. DGE profiles were generated by counting for each sample the number of unique UMIs associated with each RefSeq genes. DGE-sequenced samples were acquired from five sequencing runs. Sequenced samples with at least 50000 counts and 6000 expressed genes were retained for further analysis.

#### Transcriptomics analyses from DGE-Seq data

Samples were filtered out if the number of unique UMIs was inferior to 50000 and the number of expressed genes inferior to 6000; 165 samples passed those cutoffs. Samples were normalized using same strategy as described in the DESeq2 method (Love et al., 2014). For performing dimension reduction and clustering, samples were logged using a log2(x+1) transformation. The five different runs were merged using ComBat (Johnson et al., 2006) from the library “sva” in parametric mode, using technical replicates between batches as references for computing batch effects. Each gene expression of the corrected values was subtracted by the minimum of the gene expression before the batch correction. This step does not change the relative expression of genes; however, it permits an easier interpretation of the expression values as minimums cannot be less than zero. Finally, each set of technical replicates were merged. The resulting samples consist in the median of each gene for its set of technical replicates. A set of over-dispersed genes was determined for computing the correlation heatmap from **Fig 1C** and the PCA from **Fig3A**. To pick these, the coefficient of variation of each gene from the normalized adjusted expression was fitted by the mean expression of each gene, using a LOESS method. Genes with a positive residual for the regression were marked as over-dispersed. This leads to a total of 2770 genes. Pathway eigengenes and their gene contribution were computed with the following steps: first, the gene sets corresponding to each pathway were downloaded on the KEGG database. Pathways with at least 4 genes existing in our data were conserved. Second, a PCA was computed for each pathway, using the gene set of the pathway as the input of the PCA. The first component of each PCA was designated as “pathway eigengene”.

#### Transcriptomics analyses from single-cell RNA-Seq

Petropoulos and Zhou datasets were normalized and log transformed using scran (Lun et al., 2016), then merged with *fastMNN* from the R library “batchelor” (Haghverdi et al., 2018) with the parameter *k* set up to 420. UMAP was performed on all available genes with the R library “uwot”. The n_neighbors parameter was set to its maximum (6838), and min_dist = 0.2. A first cell clustering was done using Monocle3 (Qiu et al., 2017), then, preimplantation clusters were re-annotated using clusters found in Meistermann et al. (Meistermann et al., 2019a). First, coordinates of centroids of these previous clusters were determined on the UMAP using Petropoulos dataset, then, cells were attributed to the cluster with the nearest centroid in term of Euclidean distance. To determine the markers of each clusters, genes with at least a mean 2 normalized count in DGE-Seq and 4 normalized count in single-cell RNA-Seq were kept for computing AUROCs. Genes with an AUC of at least 0.85 were designated as markers. Markers determined by (Xiang et al., 2020) were also filtered, with a mean expression of at least 2 normalized counts in DGE-Seq data.

#### Data and software availability

Raw data is available on ENA at accession number: PRJEB34037.

**Figure S1 – related to Figure 1.**
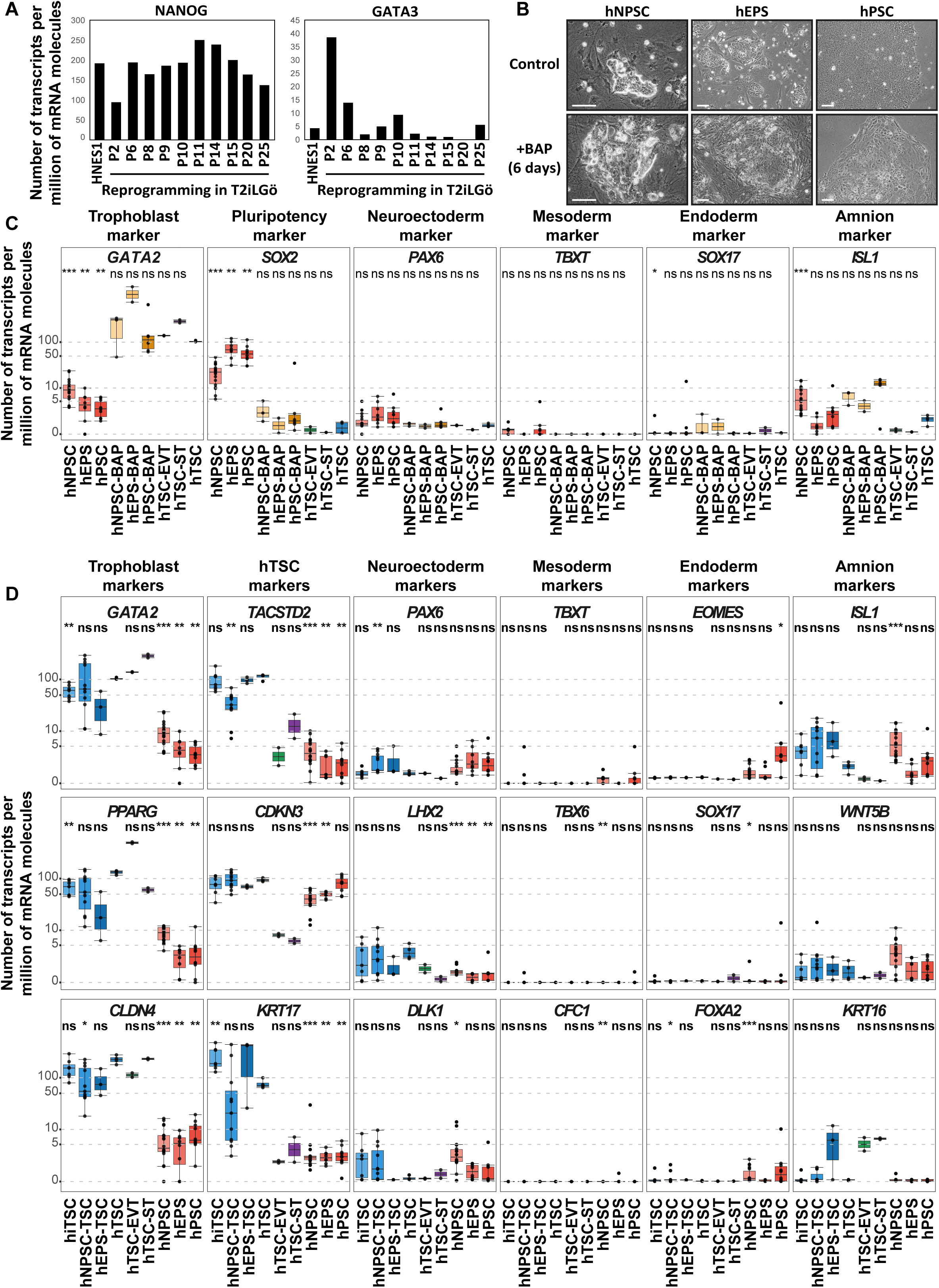
(A) Bar plots showing concomitant bulk gene expression of the pluripotency-specific transcription factor *NANOG* and the trophoblast-associated transcription factor *GATA3* in early intermediates during experiments of human somatic cell reprogramming towards naive pluripotency (P2 to P6). Absolute gene expression levels are given as number of transcripts per million of mRNA molecules. Under these culture conditions, *GATA3* expression is lost over passages while *NANOG* expression is stable. Data from (Kilens et al., 2018). (B) Phase contrast images of hNPSC, hEPS and hPSC treated with BAP (bottom). Untreated cells are shown as controls (top). (C) Gene expression levels of indicated lineage markers are shown for BAP-treated hNPSC, hEPS and hPSC along with the relative controls. Previously established hTSC and the differentiated ST and EVT are included for comparison. Expression levels are given as number of transcripts per million of mRNA molecules. A Wilcoxon-Mann-Whitney statistical test was performed, with embryo- and placenta-derived hTSC taken as the reference group. Stars indicate statistical significance of the difference: *p-value < 0.05; **p-value < 0.01; ***p-value <0.001. (D) Gene expression levels of indicated lineage markers are shown for hiTSC, hcTSC, embryo- and placenta-derived hTSC, along with hNPSC, hEPS and hPSC. The differentiated ST and EVT are included for comparison. Expression levels are given as number of transcripts per million of mRNA molecules. A Wilcoxon-Mann-Whitney statistical test was performed, with embryo- and placenta-derived hTSC taken as the reference group. Stars indicate statistical significance of the difference: *p-value < 0.05; **p-value < 0.01; ***p-value <0.001. Scale bars: 100 µm.

**Figure S2 – related to Figure 2.**
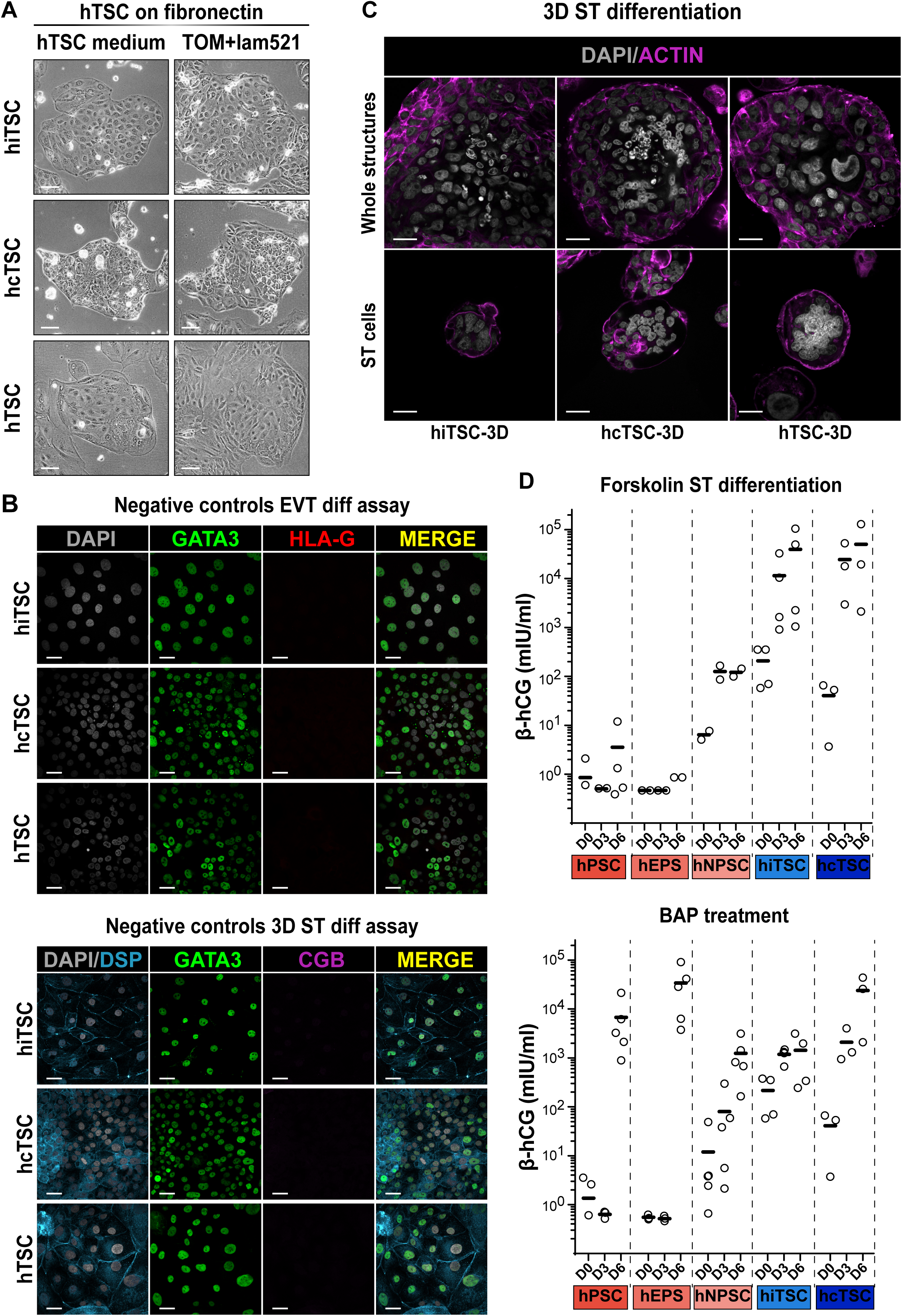
(A) Brigthfield pictures of cells prior to differentiation. Left: h(i/c)TSC were cultured on fibronectin in hTSC medium prior to EVT and ST differentiation assays. Right: h(i/c)TSC were cultured on fibronectin and laminin521 in trophoblast organoid medium (TOM) prior to 3D-ST differentiation experiments. (B) Immunofluorescence images of undifferentiated h(i/c)TSC stained for the trophoblast-associated transcription factor GATA3 and the extravillous trophoblast-specific surface marker HLA-G (top) or the syncytiotrophoblast-associated markers DSP and CGB along with GATA3 (bottom). Nuclei were stained with DAPI. (C) Immunofluorescence images of 3D-ST structures derived from hiTSC, hcTSC,and hTSC lines stained for ACTIN, highlighting the formation of syncytiotrophoblast cells (center of structures). Nuclei were stained with DAPI. (D) Scatter plots of secreted β-hCG levels in mIU/ml for indicated cell lines at day 0, 3, or 6 of ST differentiation assays. Top: results for forskolin treatment. Bottom: results for MEF-BAP treatment. Each biological replicate is represented by a dot. Bars represent averages. Scale bars (A): 100 µm; (B-C): 30 µm.

**Figure S3 – related to Figure 3.**
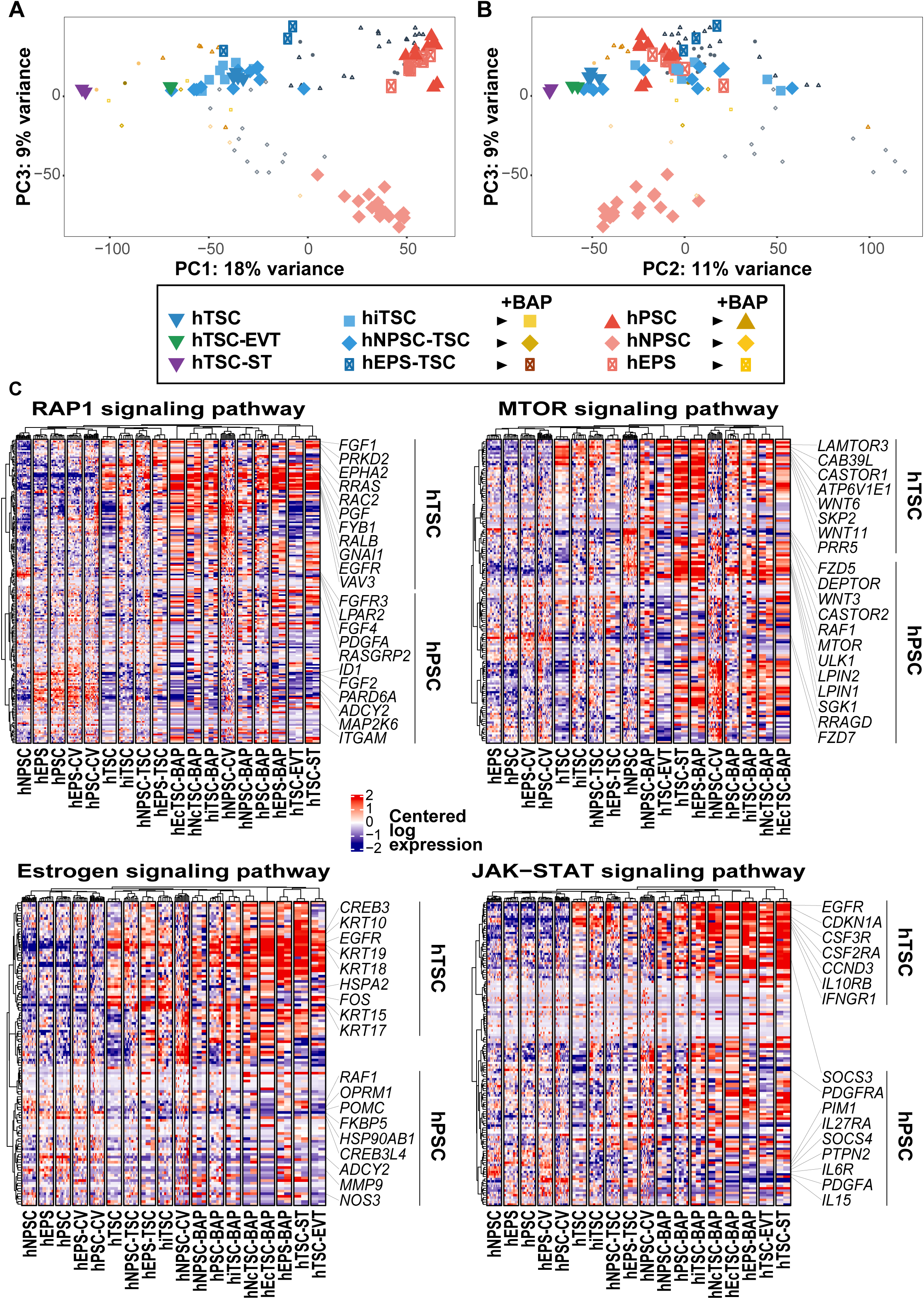
(A-B) PCA of h(i/c)TSC and hPSC in maintenance, differentiation or conversion media. PC1 vs PC3 (A) and PC2 vs PC3 (B) are presented. (C) Heatmap presentation of genes from selected signaling pathways. For each pathway, genes enriched in h(i/c)TSC or hPSC are highlighted.

**Figure S4 – related to Figure 4.**
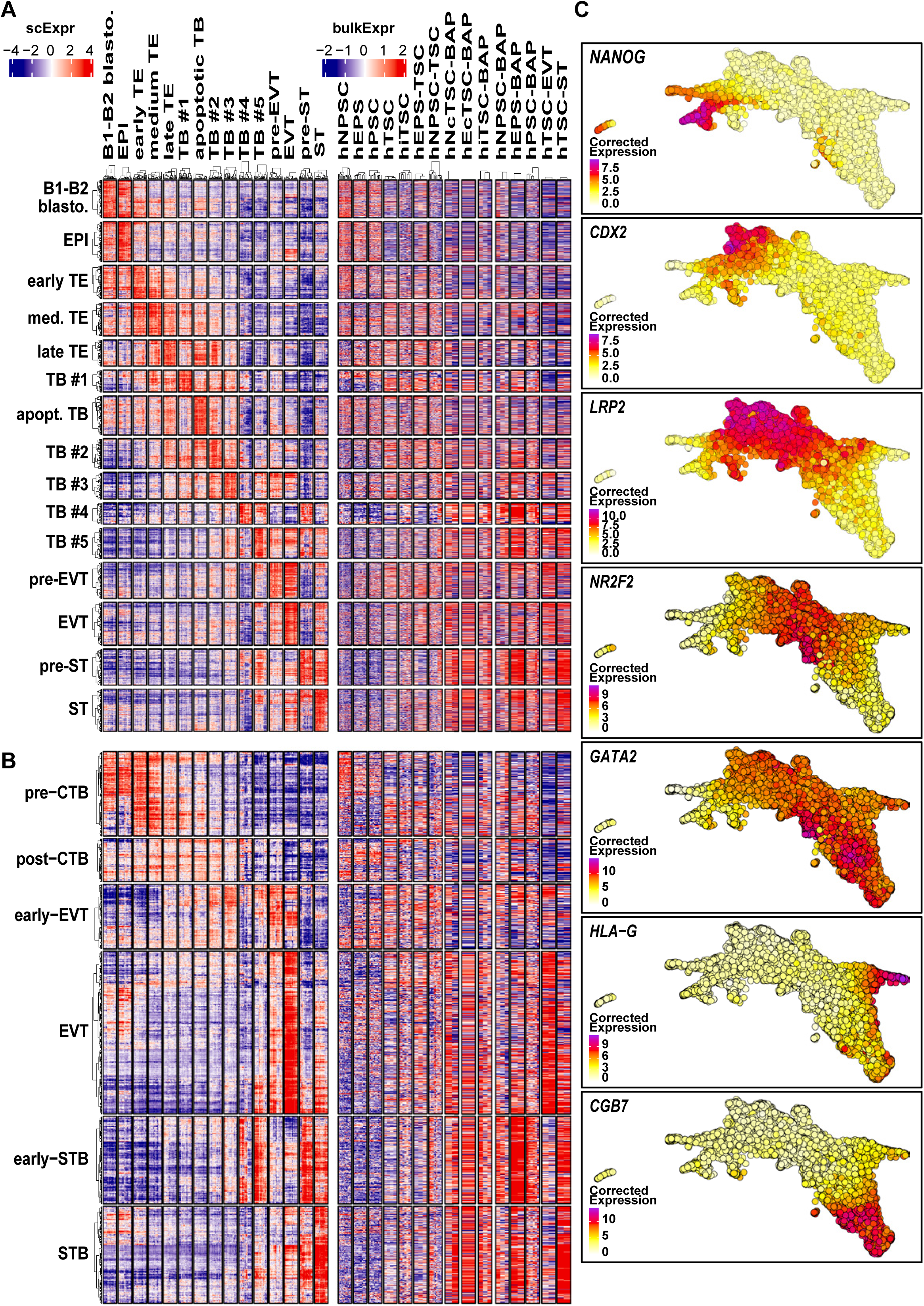
(A-B) Heatmap representation of genes associated with single-cell clusters (A) or gene sets (B, (Xiang et al., 2020)) for pre-implantation CTB, post-implantation CTB, early-EVT, EVT, early-STB and STB across scRNAseq peri-implantation embryo clusters, hPSC, h(i/c)TSC and differentiated cells. (C) Projection of expression levels for selected genes on the UMAP.

**Table S1. Description of cell lines and primer sequences**

This spreadsheet contains 2 tabs:

(A) Description of all cell lines used in this study, comprising induced, converted, embryo- and placenta-derived hTSC, along with hNPSC, hEPS and hPSC lines. Genetic background, culture conditions, and generation method are described for each cell line along with quantitative aspects of reprogramming and conversion experiments: starting number of cells, density of colonies, frequency of passages, split ratios and maximal number of passages tested.

(B) Nucleotide sequences of primers used for RT-qPCR.

**Table S2. Expression data, sample annotations and pathway component eigengene contributions**

This spreadsheet contains 7 tabs:

(A) Sample annotation for DGEseq data. The “M.” mention preceding sample names indicates technical replicates which correspond to samples that were included in two or more sequencing runs, and that were merged as one resulting sample after batch correction.

(B) Normalized expression table.

(C) Log2-transformed expression table.

(D) Expression table in UPM (units per million). Gene expression levels correspond to the number of specific transcripts per million of total mRNA molecules.

(E) Sample annotation for scRNAseq data. Developmental timing of cells is indicated in DPF (days post-fertilization). “Meistermann clusters” correspond to clusters identified in our recent report (Meistermann et al., 2019a), which focused on pre-implantation datasets. These clusters have been reused to annotate the pre-implantation clusters in this study, while additional clusters of post-implantation cells have been identified here and are regrouped as “peri-implantation clusters”. Coordinates of cells in the UMAP are indicated as UMAP D1 and UMAP D2.

(F) Comprehensive list of markers for the 19 clusters of cells identified in the peri-implantation human embryo.

(G) List of KEGG pathway component genes showing their contributions to pathway eigengenes.

## Notes

### Competing Interest Statement

Dimitri MEISTERMANN is supported by FINOX forward grant initiative.
Gael CASTEL and Laurent DAVID have a provisional patent filled on the generation of human induced trophoblast stem cells.

